# The Impact of Dance Movement Therapy on Subjective Well-Being and Quality of Life in Healthy Older Adults

**DOI:** 10.1101/2025.10.20.683423

**Authors:** Laura Sebastiani, Said Daoudagh, Giacomo Ignesti, Marina Raglianti, Paolo Paradisi

## Abstract

Undertaking physical and social activities such as dance and music have positive effects on the mental well-being, cognitive and physical abilities of older people. Dance Movement Therapy (DMT) is a complementary practice that uses dance and movement to stimulate creativity, enabling individuals to engage in positive interpersonal connections within a group. Therapeutic approaches involving DMT have been proven to promote positive mental health outcomes in various patient groups. This study aimed to evaluate the possible efficacy of DMT on the subjective perceptions of quality of life, satisfaction with life, and personal well-being of a sample of healthy adults aged over 65. Participants (n=25; females=17) took part in an 8-week DMT cycle on a once-weekly basis. The DMT cycle was preceded and followed by a one-hour session during which participants filled out the World Health Organization Quality of Life assessment (WHOQoL-BRIEF), the Satisfaction With LifeScale (SWLS), and the Wellbeing Index (PWI) scale. Comparison of pre- and post-DMT scores by repeated measures ANalysis Of VAriance (rmANOVA) showed a positive effect of DMT, especially in terms of overall quality of life and social relationships. Participants in the over-70s age group who showed significantly lower scores than their younger counterparts in terms of overall quality of life, Social subscale, and PWI, exhibited analogous DMT-related changes. Our findings suggest that DMT has the potential to foster a shared positive state of mind among older individuals, thereby facilitating social interaction not only among the elderly population but also among its most senior members.

## 1. Introduction

A substantial body of research indicates that adopting an active lifestyle, integrating physical and social activities, is the key to optimal health and general well-being in persons of all age groups Piercy et al. (2018), Sanchís-Soler, Sebastiá-Amat, and Parra-Rizo (2025) and with different abilities Biagini, Bastiani, and Sebastiani (2022). In particular, it is argued that active engagement in socio-cultural practices, such as music and dance, has the potential to enhance and maintain well-being as well as the quality of life experienced by individuals. Interestingly, the available evidence suggests that taking part in structured dance, irrespective of its specific style, is as good as other types of organised physical activity for improving mental well-being and cognitive skills Fong Yan et al. (2024). Furthermore, dance intervention exhibited a high level of adherence in comparison to aerobic and resistance exercise, therefore suggesting that structured dance constitutes an effective alternative to exercise in terms of enhancing physical function and quality of life.

In the context of the ageing process, the identification of appropriate intervention strategies could be important for the improvement of the well-being of older people, who are a particularly vulnerable population in terms of both physical and mental health. Actually, undertaking physical-social activities has been demonstrated to exert a multitude of positive effects in older persons, including the reduction of depression and disability rates Hwang and Braun (2015). In addition, social dances have been demonstrated to be a particularly efficacious means of engendering positive psychological health effects in elderly patients (i.e., QoL and internal motivation in chronic heart failure Kaltsatou, Kouidi, Anifanti, Douka, and Deligiannis (2014); mental health and quality of life in patients with Parkinson’s Disease Cheng, Quan, and Thompson (2024)).

Dance Movement Therapy (DMT) ^1^ is one of many complementary practices used to improve general well-being and is deeply rooted in the principles of embodied cognition, which posits that our cognitive processes are shaped and informed by our bodily interactions with the world de la Parra López and Panhofer (2023); Goodill (2024); Kiverstein and Miller (2015); Paradisi, Raglianti, and Sebastiani (2021); L. Shapiro and Stolz (2019); L. A. Shapiro (2019). To be more specific, DMT can be defined as a human-centred psychotherapeutic approach that utilises dance and movement to stimulate a creative process, thereby enabling individuals to establish relationships with one another within the group dynamics Dulicai and Hill (2007); Winters Fisher (2019). The group consists of individuals who interact, integrate, and relate to one another, thus forming a collective unit Overby, Shanahan, and Young (2024) (see, in particular, Goodill (2024)). It is used in a variety of situations, including one-to-one and group meetings, but most commonly in small group meetings. Therapeutic approaches involving DMT intervention have been used with various categories of patients; for example, a 10-week group DMT intervention has demonstrated efficacy in enhancing resilience, as well as mood, stress, relaxation, and pain in chronic pain patients Shim et al. (2017). In addition, improvements in cancer-related symptoms such as distress and anxiety have been reported in cancer patients participating in a virtual DMT protocol Bryl et al. (2024).

However, most studies are qualitative and lack objective measures of DMT’s effectiveness in promoting individual well-being. Furthermore, although DMT has been used in elderly patients - i.e., incorporating DMT elements into physical exercises has been shown to be effective in improving the fitness and overall functioning of elderly nursing home patients in wheelchairs Woloszyn, Wísniowska-Szurlej, Grzegorczyk, and Kwolek (2021), and weekly group dance sessions including DMT elements have improved the mood of older patients in an acute hospital setting Bungay, Hughes, Jacobs, and Zhang (2022)-there is little information available on the efficacy of DMT in healthy elderly individuals.

The aim of the present study was to assess the subjective perception of quality of life, life satisfaction, and personal well-being in a sample of healthy older people taking part in an 8-week DMT intervention. To this end, a series of validated questionnaires were administered to all participants before the start of the DMT intervention and after its completion. Based on previous results in patients, we expect that DMT may also have a positive effect on perceived quality of life and well-being in healthy older people.

**Outline** The paper is structured as follows: Section 2 illustrates the methods used in the study, including the design and participants, research instruments, procedure, and the DMT protocol employed. Section 3 presents a preliminary data analysis, whereas Section 4 reports the results of our investigation. Finally, Section 5 discusses the findings and outlines the limitations and future perspectives of the present research.

## 2. Methods

### 2.1. Design and Participants

The present study evaluated the impact of DMT on the subjective perception of the quality of life, satisfaction for life, and personal well-being in a sample of Italian people over 65 years of age. The recruitment advertisements were prepared and distributed via flyers that were left at territorial associations and gyms, and also through local newspaper advertisements. Participants who responded to the advertisement were interviewed by PP or SD to collect general personal data (e.g., age, sex, previous job, general health, prior and current motor activity) to create a database from which suitable participants could be selected. Participants were eligible for inclusion in the study if they were not affected by neurological, psychiatric, or physical disorders, were able to carry out their daily activities independently, and reported participating in leisure motor activities to a moderate extent, such as walking and cycling.

A total of 27 subjects (males=10; females=17) were found to be eligible and enrolled in the study. The recruited participants took part in the eight-week DMT cycle on a once-weekly basis. The DMT cycle was preceded (pre-DMT) and followed (post-DMT) by a session during which participants filled out a series of questionnaires. For all subjects, the pre-DMT and post-DMT sessions were completed within one week prior to and following the DMT intervention, respectively.

In accordance with the indication of the DMT professional who conducted the DMT intervention, the maximum number of participants permitted by the DMT was set at 14. This sample size was deemed appropriate for the capacity of the gym room, but primarily because smaller groups facilitate the development of group dynamics. Consequently, two eight-week DMT cycles were scheduled, and participants were assigned to one cycle or the other in order to obtain two groups (1st and 2nd cycle) of similar size and with a similar female-to-male ratio. However, one participant in the 1st cycle attended only 3 of the 8 DMT meetings and was therefore not included in the subsequent analysis. A participant of the 2nd cycle was also excluded from the analysis due to a fall from a bicycle, which resulted in impaired movement and a limited ability to interact with others during subsequent DMT meetings. 5 All other participants engaged in a minimum of seven out of eight DMT meetings and reported no motor disturbances throughout the DMT cycle. Thus, the final sample comprised two groups: 1st cycle, n=12, and 2nd cycle, n=13. Demographic data are presented in Table 1. This study was performed in accordance with the Declaration of Helsinki ethical standards and approved by the Committee on Bioethics of the University of Pisa (resolution no. 29/2024 of 26/07/2024). The participants read and signed the informed consent to participate in the study.

**Table 1.** Demographic characteristics of participants in the two intervention cycles.

|  | Cycle 1 | Cycle 2 |
| --- | --- | --- |
| **Sample size** | N=12 | N=13 |
| **Age (mean ± standard deviation)** | 68.25±3.41 | 73.67±6.17 |
| **Sex (%)** |  |  |
| female | 66.7 | 69.2 |
| male | 33.3 | 30.8 |
| **Previous Job** | N | N |
| Unemployed | 1 | 0 |
| Elementary occupations | 0 | 1 |
| Factory worker | 0 | 1 |
| Artisan profession | 0 | 2 |
| Service and sales workers | 1 | 0 |
| General Office Clerks/Secretaries | 5 | 3 |
| School and Early Childhood Teachers | 1 | 3 |
| Technicians | 2 | 1 |
| University and Higher Education Teachers | 0 | 1 |
| Managers/Directors | 2 | 1 |
| **General health (subjective evaluation)** | N | N |
| fairly good | 1 | 5 |
| good | 7 | 4 |
| very good | 3 | 4 |
| excellent | 1 | 0 |
| **Ongoing leisure motor activities** | N | N |
| walks | 10 | 13 |
| cycling | 6 | 7 |
| dance | 1 | 1 |
| gym | 1 | 2 |
| swimming | 0 | 1 |
| **Previous motor activities** | N | N |
| yes | 11 | 13 |
| no | 1 | 0 |

### 2.2. Research Instruments and Procedure

Questionnaires consisted of the Italian versions of the Geriatric Depression Scale (GDS) Galeoto et al. (2018), Pittsburgh Sleep Quality Index (PSQI) Buysse, Reynolds III, Monk, Berman, and Kupfer (1989), the World Health Organization Quality of Life assessment (WHOQoL-BRIEF) De Girolamo et al. (2000), Satisfaction With Life Scale (SWLS) Di Fabio and Gori (2016), Wellbeing Index (PWI) Verri et al. (1999),

#### 2.2.1 Geriatric Depression Scale - GDS

The GDS is a 30-item instrument designed to evaluate the emergence of depression in elderly individuals. The scale is composed of a set of questions and statements that are designed to assess the mood, feelings, and behaviour of older people. It looks at how they feel about their daily activities, their general mood, their sense of worthlessness, and any symptoms related to their memory and thinking. Users respond in a “Yes/No” format.

#### 2.2.2. Pittsburgh Sleep Quality Index - PSQI

The PSQI is a one-month self-report questionnaire that is widely used to evaluate sleep quality. The tool is considered valuable due to a multifaceted approach, incorporating both subjective experiences and objective parameters. The global PSQI score, which ranged from 0 to 21, is obtained by the sum of seven component scores. Each component score ranges from 0 to 3, with 3 indicating the greatest degree of dysfunction or disturbance. The higher the score, the poorer the sleep quality, with a score over 5 suggesting substantial sleep difficulties.

#### 2.2.3. WHOQoL-BREF

The WHOQoL-BRIEF is a quality-of-life assessment that utilizes a 26-item self-report scale and has demonstrated cross-cultural applicability. The WHOQoL-BRIEF is a tool designed to assess individuals’ psychological and physical well-being. However, it also enables subjects to offer a subjective evaluation of the role that their social environment plays in their perceived general state. In more detail, the WHOQoL-BRIEF allows the measurements of four domains associated with the quality-of-life (QoL) construct: physical health, psychological, social relationships, and environment domains, as well as overall QoL (item 1: “How would you rate your QOL?”) and general satisfaction with health (item 2: “How satisfied are you with your health?”).

The scale is assessed on a 5-point Likert scale, with higher scores denoting a higher self-perception of the QoL.

#### 2.2.4. Satisfaction With Life Scale - SWLS

The SWLS is a 5-item self-evaluation scale designed to measure global judgments of one’s life satisfaction. Participants were required to indicate on a 7-point scale how much they agree or disagree with each of the five items (from 1, strongly disagree, to 7, strongly agree).

#### 2.2.5. Wellbeing Index - PWI

The PWI scale is a self-evaluation scale that provides a domain-level representation of subjective global life satisfaction. The core questionnaire consists of 7 questions designed to assess satisfaction with specific life domains: standard of living, personal health, achievement in life, personal relationships, personal safety, community connectedness, and future security. In addition to these core items, the questionnaire contains two optional questions regarding general life satisfaction and spiritual or religious satisfaction. The scores of the core set of items plus the religion item, if answered, are averaged to produce a Personal Wellbeing Index. The item relative to satisfaction with life as a whole is analyzed as a separate variable and generally used to test the construct validity of the PWI. The participants were required to indicate on a 10-point Likert scale how much satisfied they are (from 0, very unsatisfied, to 10, very satisfied).

### 2.3. DMT Protocol

The experimental protocol consisted of eight weekly DMT meetings lasting approximately one hour each and taking place in a standard gym facility with sufficient space for unrestrained movement. As a DMT professional (conductor) associated with APID APID (2025), MR conducted the DMT meetings. For each meeting, the general outline of the settings, including suggestions, possible tools (e.g., laces) and the sequence of musical pieces were previously planned. In the 2nd cycle of DMT, the same settings were applied, and the order of the meetings was maintained as in the 1st cycle.

Each meeting began with a “welcome” phase. This was conducted with participants taking a position within a circle or, in one or two meetings, along a row, under the guidance of the conductor. The aim of the *welcome* phase was to allow participants to introduce themselves by having them say their names and then calling each other by name, in order to promote familiarisation between the group members and provide an opportunity to get to know each other better. The subsequent phase comprised a “warm-up”, accompanied by music, during which participants initiated preparatory activities for their bodies, including articulation of the joints and mobilisation of various anatomical structures. The core of each DMT meeting was centred around a specific thematic element (e.g., exploration of the space around the body; finding unusual ways to move). The theme was typically developed within three phases that typically involved individual, dyadic, and group motor dialogues, with or without the use of accessories (e.g., elastic ribbons, masks, veils). All phases were accompanied by appropriate music. The concluding segment of the DMT meeting typically culminated in a group activity that was structured around a soundtrack characterised by pleasant and/or joyful music. In some meetings, the conductor proposed a short “final greeting” phase with all participants standing in a circle, hand in hand, all together taking two steps to the right and one to the left.

## 3. Data Analysis

### Preliminary analysis

The normality of data distribution was evaluated for each scale using the Kolmogorov-Smirnov test. As all the scales exhibited normal distribution, parametric statistical analyses were employed.

For each scale, the internal consistency was assessed by means of Cronbach’s *α*. For scales that have fewer than ten items, an *α* value greater than 0.5 is considered acceptable. Cronbach’s *α* for PWI, SWLS and WHOQoL were, respectively, 0.881, 0.846, 0.817 in the pre-DMT session and 0.916, 0.878, 0.756 in the post-DMT session. The group’s mean GDS scores indicated an overall absence of depressive symptoms (mean ± Standard Deviation (SD), 7.68 ± 4.27), with only nine out of 25 participants reporting scores above the cutoff (cutoff = 9) consistent with mild depression Galeoto et al. (2018). No changes in GDS scores occurred following the DMT intervention (7.26 ± 4.84). The group’s mean PSQI scores (mean ± SD, 5.20 ± 2.83) indicated a very slight alteration in sleep quality (cutoff = 5) which is consistent with the agerelated physiological changes in some sleep parameters (e.g., reduction in total sleep time; increase in night awakenings) Crowley (2011). Similar PSQI scores were found following the DMT intervention (5.25 ± 2.88).

Preliminary repeated measure ANalysis Of VAriance (rmANOVA) performed on all the scales (WHOQol subscales, SWLS, PWI) with DMT as within-subject factor (pre-DMT, post-DMT) and Cycle as between-subject factor (Cycle 1 and 2) did not reveal any Cycle effect (MANOVA (F(7,17)=1.395, p=0.27) nor DMT x Cycle interaction (F(5,20)=1, 915, p=0.13). Therefore, for successive analysis, data from Cycle 1 and 2 were collapsed together.

### Statistical Design

For all the questionnaires/scales we compared the mean scores of the two sessions by means of rmANOVA, with DMT (pre-DMT and post-DMT) as a within-subjects factor, and Age (65-69 and over 70) and Sex (female, male) as between-subject factors. Significance was set at *p <* 0.05. For univariate comparisons, Bonferroni correction for multiple comparisons was applied. The SPSS.15 statistical package was used for all analyses.

### DMT evaluation sheets

Following the completion of each DMT cycle, the conductor provided a qualitative assessment of the group’s performance using the evaluation form typically used by DMT professionals according to APID guidelines APID (2025) The evaluation took into account several aspects, including the group’s attitude towards each other and towards the leader, their creative ability, how they used the space, the types of movements they performed, and the body parts they used.

## RESULTS

### Questionnaires results

The scores (mean + SD) reported by the two groups of age in the pre-DMT and post-DMT sessions are shown in Table 2.

**Table 2.** Questionnaires’ Results.

| Questionnaire | Age | Session | Mean | SD | Confidence Interval Limits |  |
| --- | --- | --- | --- | --- | --- | --- |
|  |  |  |  |  | Lower | Upper |
| Overall QoL | 65–69 | pre | 73.21 | 8.29 | 68.09 | 78.33 |
|  |  | post | 77.68 | 11.16 | 71.82 | 83.54 |
| | $\geq 70$ | pre | 63.64 | 10.39 | 57.86 | 69.41 |
|  |  | post | 71.59 | 9.82 | 64.98 | 78.20 |
| Physical Health | 65–69 | pre | 77.81 | 9.20 | 71.91 | 83.70 |
|  |  | post | 79.34 | 11.81 | 72.70 | 85.97 |
| | $\geq 70$ | pre | 69.16 | 12.30 | 62.51 | 75.80 |
|  |  | post | 72.08 | 12.25 | 64.59 | 79.57 |
| Psychological | 65–69 | pre | 67.56 | 14.17 | 60.13 | 74.98 |
|  |  | post | 68.15 | 11.96 | 61.70 | 74.61 |
| | $\geq 70$ | pre | 59.85 | 12.40 | 51.47 | 68.22 |
|  |  | post | 55.83 | 11.29 | 48.55 | 63.12 |
| Social Relationships | 65–69 | pre | 71.55 | 15.76 | 62.76 | 80.34 |
|  |  | post | 77.08 | 12.63 | 70.60 | 83.56 |
| | $\geq 70$ | pre | 48.03 | 16.09 | 38.11 | 57.95 |
|  |  | post | 60.98 | 10.42 | 53.67 | 68.30 |
| Environment | 65–69 | pre | 68.97 | 12.35 | 61.43 | 76.52 |
|  |  | post | 72.77 | 9.80 | 66.18 | 79.36 |
| | $\geq 70$ | pre | 63.07 | 15.17 | 54.56 | 71.58 |
|  |  | post | 62.50 | 14.18 | 55.07 | 69.94 |
| PWI | 65–69 | pre | 78.66 | 7.35 | 72.31 | 85.00 |
|  |  | post | 81.16 | 6.91 | 76.03 | 86.29 |
| | $\geq 70$ | pre | 66.02 | 15.26 | 58.86 | 73.18 |
|  |  | post | 64.99 | 11.65 | 59.20 | 70.77 |
| SWLS | 65–69 | pre | 25.07 | 5.24 | 22.18 | 27.96 |
|  |  | post | 26.29 | 4.79 | 23.84 | 28.74 |
| | $\geq 70$ | pre | 21.27 | 5.20 | 18.02 | 24.53 |
|  |  | post | 21.09 | 3.91 | 18.33 | 23.86 |

Repeated measure ANOVA yielded a significant DMT effect (MANOVA, F(7, 15) = 3.777, p = 0.015, *η*2 = 0.638, power=0.875). In fact, as can be observed in Figure 1, which shows the mean scores of the various questionnaires in the pre-DMT and post-DMT sessions, the post-DMT scores tend to be higher than the pre-DMT ones in all the scales. However, univariate comparisons yielded a significant DMT effect for the general QoL (F(1,21)=10.861, p=0.003, η2=0.342, power=0.881) and the WHOQoL Social relationships subscale (F(1,21)=11.117, p=0.003, η2=0.346, power=0.888), only. No significant interactions between DMT and Age were found for any of the WHOQoL dimensions, nor for the SWLS and PWI scales. In particular, as shown in Table 2, an increase in overall QoL and Social relationship dimension was found in the two groups of age.

**Figure 1.**
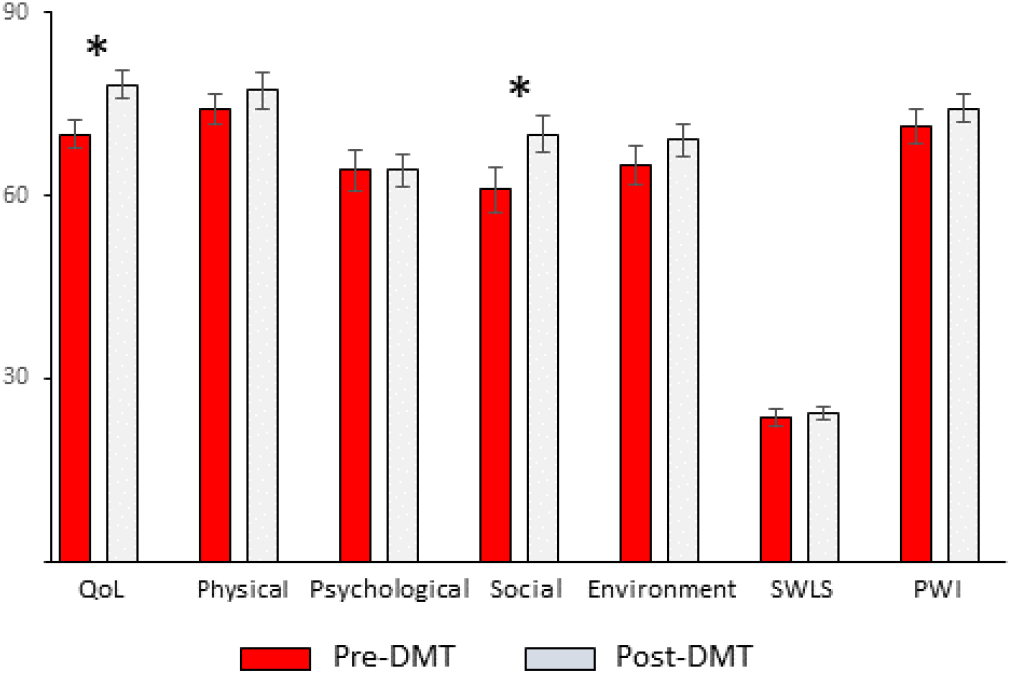
The scores (mean ± standard error of the mean(SEM)) obtained in the WHOQoL (overall QoL and Physical Health, Psychological, Social relationships, Environment domains), SWSL, and PWI scales in the pre-DMT (red bars) and post-DMT (light grey bars) sessions are shown. Significant differences are indicated with *∗* ( *p <* 0.007).

Univariate comparisons also yielded a significant Age effect for the general QoL (F(1,21) = 10.975, p=0.003, η2=0.343, power=0.885), WHOQoL Social subscale (F(1,21)=11.750, p=0.003, η2=0.359, power=0.904) and PWI (F(1,21)=10.574, p = 0.004, η2=0.335, power=0.873). As shown in Figure 2 and Table 2, the scores of the *65-69* group tend to be higher than the *over 70* group in all the scales. However, only the scores obtained in QoL, Social subscale, and PWI were significantly different. No Sex effects nor significant interactions between DMT, Age, and Sex were found.

**Figure 2.**
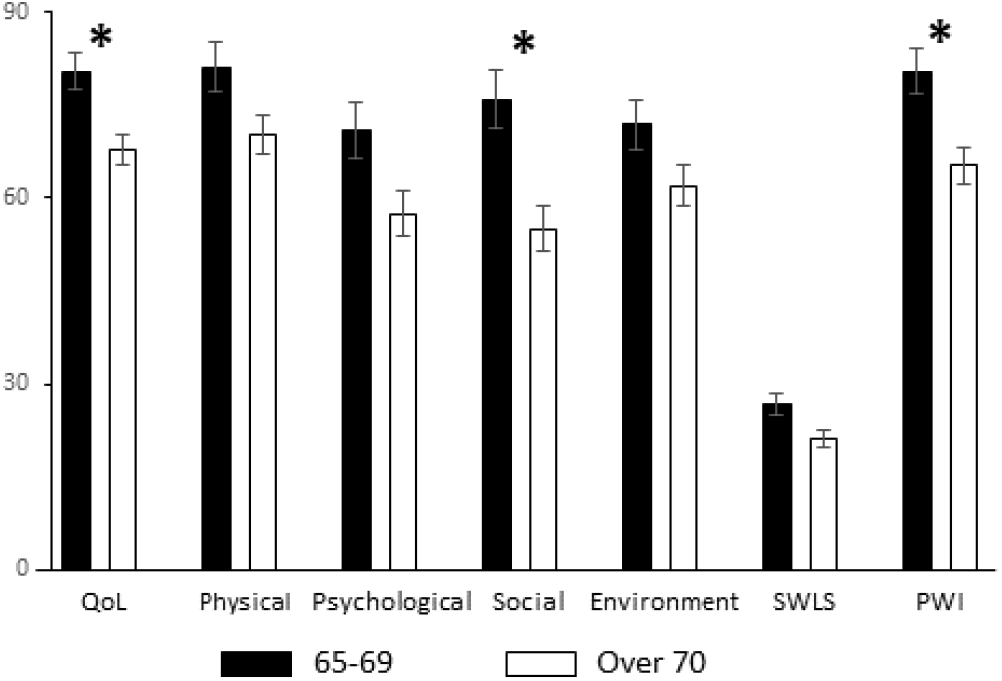
The scores (mean ± SEM) obtained in the WHOQoL (overall QoL and Physical Health, Psychological, Social relationships, Environment domains), SWSL and PWI scales in the *65-69* (black bars) and over 70 (white bars) groups are shown. Significant differences are indicated with *∗* (*p <* 0.007).

### DMT evaluation results

The conductor’s qualitative evaluation of the group’s performance evolution across the DMT intervention revealed general improvement in several areas. In fact, all participants rediscovered the pleasure of movement, experienced improvements in their motor creativity and body image, and gained better access to imaginative activities. Furthermore, the conductor highlighted the progressive emergence of a group identity through the strengthening of relationships among participants. The conductor further emphasised the presence of an atmosphere that was positive, entertaining, and engaging.

## Discussion

The present study has investigated the impact of an 8-week DMT intervention on the subjective perception of the quality of life, satisfaction for life, and personal well-being of a sample of Italian people over 65 of age. The results indicated a generally positive effect of DMT on quality of life and well-being, with significant differences in perceived overall QoL and Social relationships dimension, which suggests that partaking in the DMT protocol has helped participants to engage in positive interpersonal connections. The oldest participants (over 70) showed generally lower scores than participants under 70 in all the scales. However, both age groups exhibited analogous changes in overall Qol and the Social relationships dimension following the DMT intervention.

Regarding age and well-being, a considerable body of research has been dedicated to investigating their inter-relationship. A U-shaped relationship between age and well-being has been observed in high-income English-speaking countries. The lowest point in this relationship is around the age of 50. However, it should be noted that this pattern does not apply consistently across all populations. Indeed, a progressive decline in well-being with age has been observed in other countries, with minimal changes in the overall trend being noted Steptoe, Deaton, and Stone (2015).

Regarding the Social relationships dimension, we can make the following observations. Firstly, each DMT meeting comprised various phases characterized by different levels of complexity (i.e., the initial warm-up was conducted on an individual basis while the successive phases could involve dyadic or group interactions and/or include the use of accessories such as elastic tapes) and emotional engagement. Despite the incorporation of protracted and rhythmic pieces of music (e.g., a 15-minute excerpt from Ravel-Bolero ^2^) or settings that demanded elevated degrees of emotional involvement (e.g., the neutral mask setting), it is noteworthy that the meetings culminated in a group activity phase, typically accompanied by music of a pleasant and joyful nature, such as a waltz. Participants generally experienced this final phase as a form of emotional decompression, which typically induced happy feelings. In fact, at the end of each meeting, a general positive mood was reported by all the participants.

In addition, it could be argued that the social aspect of DMT fostered a sense of “community”, which further motivated the engagement throughout the entire DMT meeting. It is noteworthy, in fact, that all participants attended at least seven out of eight meetings. This high level of attendance, which is crucial for the intervention to be effective, could have also been facilitated by the lack of specific requirements in terms of previous experience in dance or physical activities and of a high level of fitness, which is unlikely, especially among the oldest participants. In fact, DMT does not incorporate any structured motor/dance activity; rather, it stimulates the creativity of participants, who are thus free to move in accordance with their motor possibilities, following the music in a way they choose. Furthermore, the creativity component of the programme encouraged spontaneous interaction between the participants, who were thus able to develop their own dyadic or group choreography on the spot.

Again, in terms of social interactions, our real-time qualitative observations during DMT meetings highlight mimicry episodes, particularly within the dyadic condition. This suggests a synchrony-based mechanism of interpersonal influence. We further observed that participants may also adopt different roles and coordinate their behaviour in a complementary manner. This behaviour was also evident in the group dynamics, where a leader would typically propose a new movement that could either be shared (mimicry) or developed collaboratively by the group. All these non-verbal behaviours are an indispensable component of social interactions. A plethora of studies have demonstrated that a more direct orientation towards an interaction partner and a reduction in interpersonal distance are linked to a range of prosocial outcomes (e.g., greater affiliation and cooperation; positive mood) Hove and Risen (2009); Tschacher, Rees, and Ramseyer (2014); Valdesolo, Ouyang, and DeSteno (2010). Previous studies have identified synchrony, defined as the temporal alignment of one’s movements with those of others, as a significant marker for elevated levels of social interaction Lee, Launay, and Stewart (2020). Indeed, a considerable body of research has highlighted the automatic emergence of interpersonal movement synchrony when participants perform a task together, even in the absence of explicit instruction Valdesolo et al. (2010).

In a nutshell, our findings suggest that, among other complementary practices, DMT has the potential to foster a shared positive state of mind among older individuals, thereby facilitating social interaction not only among the elderly population but also among its most senior members. This aspect is particularly relevant since it has been observed that older adults may exhibit a heightened vulnerability to social isolation and loneliness Courtin and Knapp (2017), which have been linked to a range of health concerns, such as depressive symptoms Santini et al. (2020), cognitive decline Lara et al. (2019), cardiovascular disease Valtorta, Kanaan, Gilbody, and Hanratty (2018) and an elevated risk of mortality Elovainio et al. (2017).

Our study was conducted on a sample of healthy aged people, while previous approaches involving DMT have been largely directed towards groups of patients of different ages affected by various pathologies in different domains (e.g., neurological, psychological, or psychiatric). These studies have suggested that DMT could be considered a useful complementary therapy, as it has been found to promote positive mental health outcomes, such as resilience or a reduction in distress symptoms, in these patients Bryl et al. (2024); Shim et al. (2017). The present study, conducted on a group of healthy, physically active older people, thus extends the positive effects on the psychological health of a DMT intervention to the healthy elderly population.

### Limitations and Future Perspectives

One limitation of the present study is its small sample size. However, DMT professionals strongly recommended recruiting no more than 14 participants in each DMT cycle to enable better group dynamics. Furthermore, despite a high number of people responding to the advertisement, only a few of them met the criteria for being recruited into the study. Further recruitment initiatives are underway to facilitate the scheduling of additional DMT cycles.

A further limitation is evidenced by the imbalance in the number of males and females: two-thirds of the subjects were, in fact, female. The gender imbalance in dance-related interventions for older adults has been previously reported Hwang and Braun (2015). Indeed, the majority of studies included a greater proportion of female participants, with some studies incorporating exclusively female subjects. In the present study, an attempt was made to mitigate this dishomogeneity by maintaining a comparable female-to-male ratio in both DMT cycles.

Ultimately, the study only assessed the short-term impact of DMT on well-being and quality of life. Future studies may include longitudinal assessments in both the short and long term.

## Acknowledgments

We greatly acknowledge the excellent technical assistance of Paolo Orsini.

## CRediT Roles (Authors Contribution)

LS: Conceptualization of experimental protocol; Conceptualization of data analysis; Data collection; Data curation; Formal analysis; Investigation; Methodology; Resources; Supervision; Validation; Visualization; Roles/Writing - original draft; and Writing - review & editing.

SD: Conceptualization of experimental protocol; Selection criteria and recruitment of participants; Data collection; Investigation; Visualization; Roles/Writing - review & editing.

GI: Data collection

MR: Experimental protocol; Preparation and execution of DMT settings; Data collection; Formal analysis.

PP: Conceptualization of experimental protocol; Preparation of DMT settings; Selection criteria and recruitment of participants; Data collection; Formal analysis; Funding acquisition; Investigation; Methodology; Project administration; Resources; Supervision; Writing - review & editing.

## Footnotes

1 DMT is a complementary therapy with recognized professionals in accordance with current regulations. Defined by the American Dance Therapy Association (ADTA) ADTA (2025) and the European Association Dance Movement Therapy (EADMT) EDMT (2025), In Italy, DMT professionals are associated with the Italian Professional Association of Dance-Movement-Therapy (Associazione Professionale Italiana DMT, APID APID (2025)). DMT professionals get a DMT practitioner certificate after a three-year professionalizing course with a final thesis/dissertation and a practical test.

2 The version of Ravel-Bolero we have used can be found at the following link: https://youtu.be/r30D3SW4OVw?si=dBLoN1TxlmhuCgW

